# Analysis of auxin responses in the fern *Ceratopteris richardii* identifies tissue ontogeny as a major determinant for response properties

**DOI:** 10.1101/2024.05.03.592339

**Authors:** Sjoerd Woudenberg, Melissa Dipp Alvarez, Juriaan Rienstra, Victor Levitsky, Victoria Mironova, Enrico Scarpella, Andre Kuhn, Dolf Weijers

## Abstract

The auxin signalling molecule regulates a range of plant growth and developmental processes. The core transcriptional machinery responsible for auxin-mediated responses is conserved across all land plants. Genetic, physiological and molecular exploration in bryophyte and angiosperm model species have shown both qualitative and quantitative differences in auxin responses. Given the highly divergent ontogeny of the dominant gametophyte (bryophytes) and sporophyte (angiosperms) generations, however, it is unclear whether such differences derive from distinct phylogeny or ontogeny. Here, we address this question by comparing a range of physiological, developmental and molecular responses to auxin in both generations of the model fern *Ceratopteris richardii*. We find that auxin response in Ceratopteris gametophytes closely resembles that of a thalloid bryophyte, whereas the sporophyte mimics auxin response in flowering plants. This resemblance manifests both at phenotypic and transcriptional level. Furthermore, we show that disrupting auxin transport can lead to ectopic sporophyte induction on the gametophyte, suggesting a role for auxin in the alternation of generations. Our study thus identifies ontogeny, rather than phylogeny, as a major determinant of auxin response properties in land plants.

**Summary statement:** Studies in angiosperms and bryophytes have left unresolved the roles of tissue ontogeny and species phylogeny in auxin response. We address that problem by characterizing auxin response in a fern.

## Introduction

Throughout evolution, plants adopted hormone signalling pathways to control their development, and their responses to external stimuli. Many of these phytohormones are broadly distributed, and responses are in many cases conserved among all land plant taxa, with components of land plant pathways found even in algal sisters (reviewed in (Blázquez et al., 2020; Bowman et al., 2019; Depuydt and Hardtke, 2011; Menand et al., 2007; Wang et al., 2015)). Though the initial description of such phytohormone responses and pathways was restricted to angiosperms, notably *Arabidopsis thaliana* (hereafter Arabidopsis) (e.g. auxin (Evans et al., 1994; Sieburth, 1999)), recent years have seen increased exploration in bryophytes, such as *Physcomitrium patens* (hereafter Physcomitrium) (Sakakibara et al., 2003; Thelander et al., 2018) and *Marchantia polymorpha* (hereafter *Marchantia*) (Eklund et al., 2015; Flores-Sandoval et al., 2015; Kato et al., 2015). Responses to phytohormones have thus been recorded in a variety of land plants, but these can hardly be compared because of the divergent morphologies between the different land plant clades. It is therefore difficult to define a clear evolutionary scenario for the hormonal control of plant growth and development. This difficulty is most prominent between bryophytes and tracheophytes, as their dominant generations do not share direct tissue homologies (Harrison, 2017).

Auxin, one of the most-studied and best-understood plant hormones, is crucial for many processes in bryophytes and tracheophytes (reviewed in (Kato et al., 2018)). The pathway mediating transcriptional auxin responses is conserved among all land plants and partly present in streptophyte algae (Carrillo-Carrasco et al., 2023; Mutte et al., 2018). Auxin controls many different developmental processes, including rhizoid formation in the bryophytes Marchantia and Physcomitrium (Flores-Sandoval et al., 2015; Sakakibara et al., 2003; Thelander et al., 2018) and root branching at the expense of root elongation in Arabidopsis (Casimiro et al., 2001; Dubrovsky et al., 2008; Lavenus et al., 2013). However, because of the lack of direct tissue homology, it is hard to conclude whether those processes are comparable. One exception might exist, as both Arabidopsis root hairs and Marchantia rhizoids — transcriptionally homologues structures (Jones and Dolan, 2012; Menand et al., 2007) — extend and initiate upon auxin treatment (Flores-Sandoval et al., 2015). Other processes are superficially similar, like gravitropism (Lobachevska et al., 2022; Rashotte et al., 2000; Zhang et al., 2019), but are not comparable on a cellular and tissue level. Many auxin responses in bryophytes can not thus directly be compared to those in tracheophytes, and therefore the question remains which part of the response is dependent on tissue ontogeny and which on species phylogeny.

Ferns, as sister clade to flowering plants, are positioned phylogenetically intermediate between the model bryophytes Marchantia and Physcomitrium and the model tracheophyte Arabidopsis (Donoghue et al., 2021; Puttick et al., 2018). Most important, fern lifestyles closely resemble both (thalloid) bryophytes and tracheophytes (Conway and Di Stilio, 2020). During land plant evolution, a major transition took place from a dominant haploid gametophyte generation in bryophytes to a dominant diploid sporophyte in tracheophytes. In ferns (and lycophytes), both these generations are autotrophic and free-living. From a haploid spore, a gametophytic prothallus forms that develops rhizoids, archegonia and antheridia whereas from the diploid embryo, a sporophytic plant develops with leaves and root-hair carrying roots, both innervated by vasculature. There are sparse descriptions of auxin responses in the model fern *Ceratopteris richardii* (hereafter Ceratopteris), but no clear overview of responses in both generations has been reported. Auxin treatments on Ceratopteris gametophytes are known to affect sexual differentiation, spore germination, rhizoid development, and the positioning and development of the lateral notch meristem (Chauhan, 2024; Gregorich and Fisher, 2006; Hickok and Kiriluk, 1984; Withers et al., 2023). In gametophytes of other fern species, reported effects also include cell expansion and elongation (Miller, 1961; Miller and Miller, 1965) and antheridium differentiation (Ohishi et al., 2021). Sporophytic roots from Ceratopteris show reduced growth rates upon auxin treatments and an increase in adventitious rooting, whereas no effects on lateral root initiation has been reported (Hou et al., 2004; Yu et al., 2020). In sporophytes of other fern species, cell wall extensibility and cell elongation was increased upon auxin treatment in the rachis (Cookson and Osborne, 1979), whereas — in contrast to flowering plants —auxin was reported not to promote vascular differentiation (Ma and Steeves, 1992).

Here, we mapped auxin responses in both generations of the fern Ceratopteris to uncouple tissue ontogeny from species phylogeny as determinants of auxin responses. We find that phenotypically, Ceratopteris gametophytes respond similar to Marchantia gametophyte whereas Ceratopteris sporophytes closely resemble responses of Arabidopsis sporophytes. This separation in response is further shown in their profiles of auxin-dependent gene regulation. We identify that differences in auxin-dependent gene activation are likely caused by increased levels of Aux/IAA repressor proteins in the sporophyte. Lastly, we find that auxin may control the alternation of generations. Together, our data identifies tissue ontogeny, rather than species phylogeny, as a major driver for divergence in auxin response in land plants.

## Results

### A thalloid liverwort-like response in the Ceratopteris gametophyte

To explore the growth responses to externally applied auxin in the Ceratopteris gametophyte, spores were placed on media containing different concentrations of the natural auxin Indole 3-Acetic Acid (IAA) or the synthetic auxin 1-Naphthyl Acetic Acid (NAA). Gametophytes grown prior to sexual maturation showed a clear reduction in growth in response to both auxins at all concentrations tested, except at 100 nM IAA, which appeared to slightly promote thallus growth (Fig. 1A-C). Under standard conditions, rhizoids only develop close to the spore coat (Conway and Di Stilio, 2020). With NAA however, rhizoids developed ectopically on the thallus border (Fig. S1A). Given the strong growth-inhibiting effect of both auxins on the thallus, it is difficult to disentangle potentially independent effects on growth and rhizoid formation. We therefore transferred sexual immature gametophytes to auxin-supplemented media following initial growth on control media. Upon transfer, a small growth reduction was visible but also a clear induction of (ectopic) rhizoids (Fig. 1D; Fig. S1). This response closely resembles the phenotype in Marchantia in qualitative terms (Fig. 1E). Detailed microscopic analysis revealed that the morphology of the prothallus also changed in auxin-treated gametophytes, manifested in disorganization of the meristematic notch (Fig. 1F; Fig. S1A). Most importantl, not only did we observe these morphological defects in gametophytes treated with external auxin, but also when inhibiting IAA transport with 1-N-Naphtylphtalamic Acid (NPA; Fig. 1F; Fig. S1A). The connection of auxin action with the development of the lateral meristem is in agreement with previously reported data (Withers et al., 2023) and with phenotypes reported in Marchantia (Flores-Sandoval et al., 2015). The product of the Ceratopteris lateral meristems is the female sexual organ, which establishes the first major 3D axis and will form the future sporophyte upon fertilization of its egg cell (Conway and Di Stilio, 2020). We did not observe obvious defects in the morphology of the archegonia even upon NPA treatment (Fig. S1B). Therefore, Ceratopteris gametophytes show auxin responses that are similar to those observed in gametophytes of the thalloid liverwort Marchantia.

**Figure 1:**
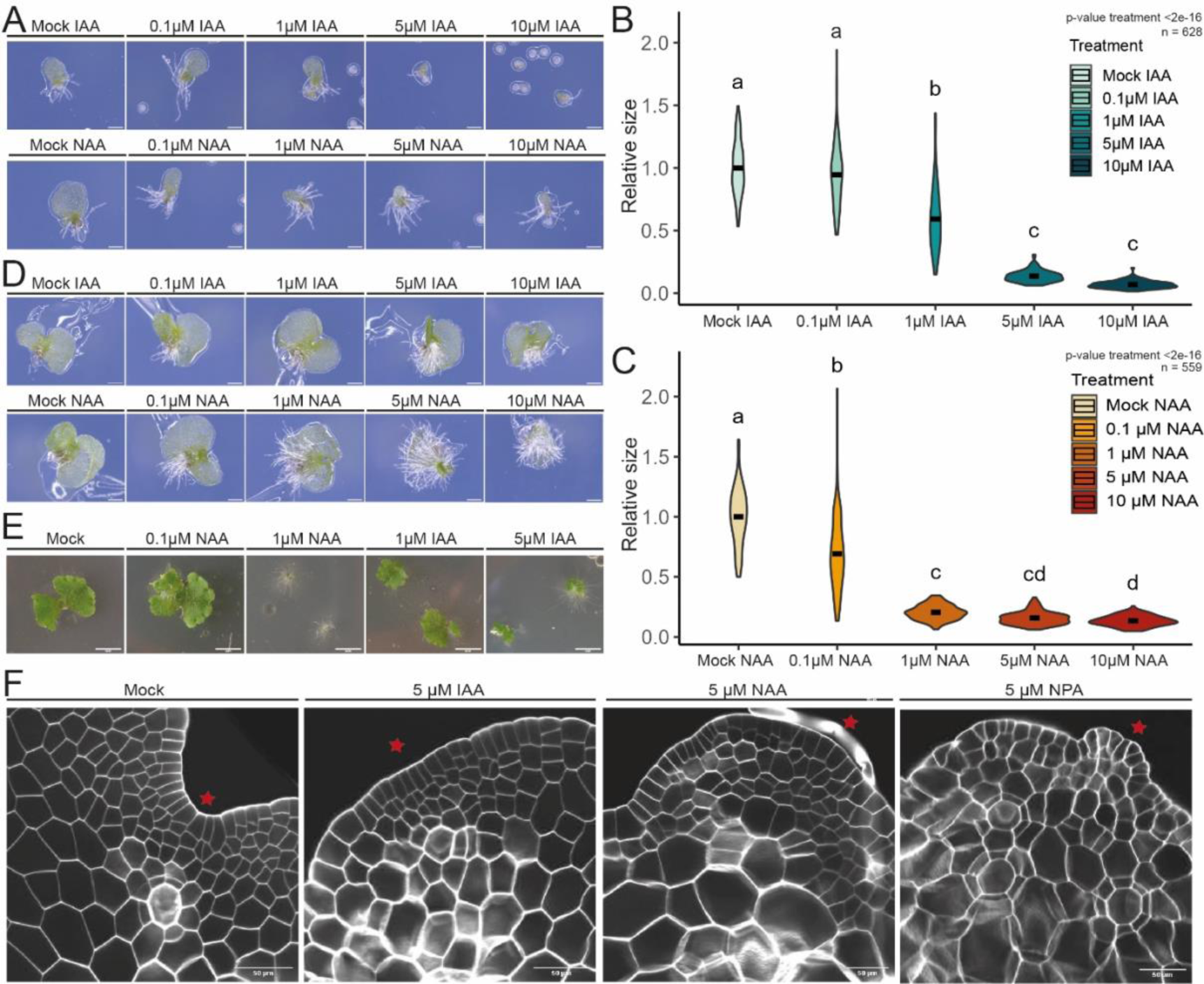
Auxin response in Ceratopteris gametophytes. *A) Gametophytes developed from spores germinated and grown on auxin-supplemented medium (at the indicated concentrations of IAA or NAA) until sexual maturity. B) Violin plots of the relative thallus size (normalized to mock treatment) upon IAA treatment or C) NAA treatment of 4 pooled replicate experiments (n>100 gametophytes per treatment). D) Gametophytes four days after transfer to auxin-supplemented medium after germination and developing a lateral meristem on unsupplemented media. E) Twelve-day-old* Marchantia *polymorpha gemmalings grown on medium supplemented with the indicated concentrations of NAA or IAA. F) Cellular organization of lateral notch meristems in thallus derived from spores transferred to auxin supplemented medium after 4 days of germination on hormone-free medium. Scale bars are 0.25 mm in A, 0.5 mm in D, 50 mm in E and 50 µm in F. Statistical significance in B and C is shown by letters based on ANOVA and Tukey pair-wise comparison (P<0.05). ANOVA P-values plotted for treatments effects in plots*.

### The Ceratopteris sporophyte shows a flowering plant-like auxin response

To test the responses of Ceratopteris sporophytes to external auxin, we transferred young (embryonic) sporophytes to medium containing different concentrations of IAA and NAA. We initially focused on the growth and branching of the root system. The ‘’primary’’ first appearing root showed a clear reduction in root growth at all IAA and NAA concentrations (Fig. 2A,B), in line with an earlier report (Hou et al., 2004). One difficulty with Ceratopteris is the lack of a true primary root. Therefore, responses were also measured on adventitious roots that are hard to perfectly synchronize and that showed a similar decrease in root growth upon auxin treatment (Fig. 2D). Interestingly, these roots also showed a clear increase of lateral root density upon auxin treatment, which is in contrast to earlier observations (Hou et al. (2004). This increased branching was pronounced after 12 days of growth in auxin supplemented medium (Fig. 2C,E), but an initial response was already visible after three days (Fig. S2A,B). Interestingly, the increased branching was not visible in the first appearing root of the young sporophytes, clearly showing that branching is root age/stage-dependent, and potentially explaining why this response is sometimes not visible.

**Figure 2:**
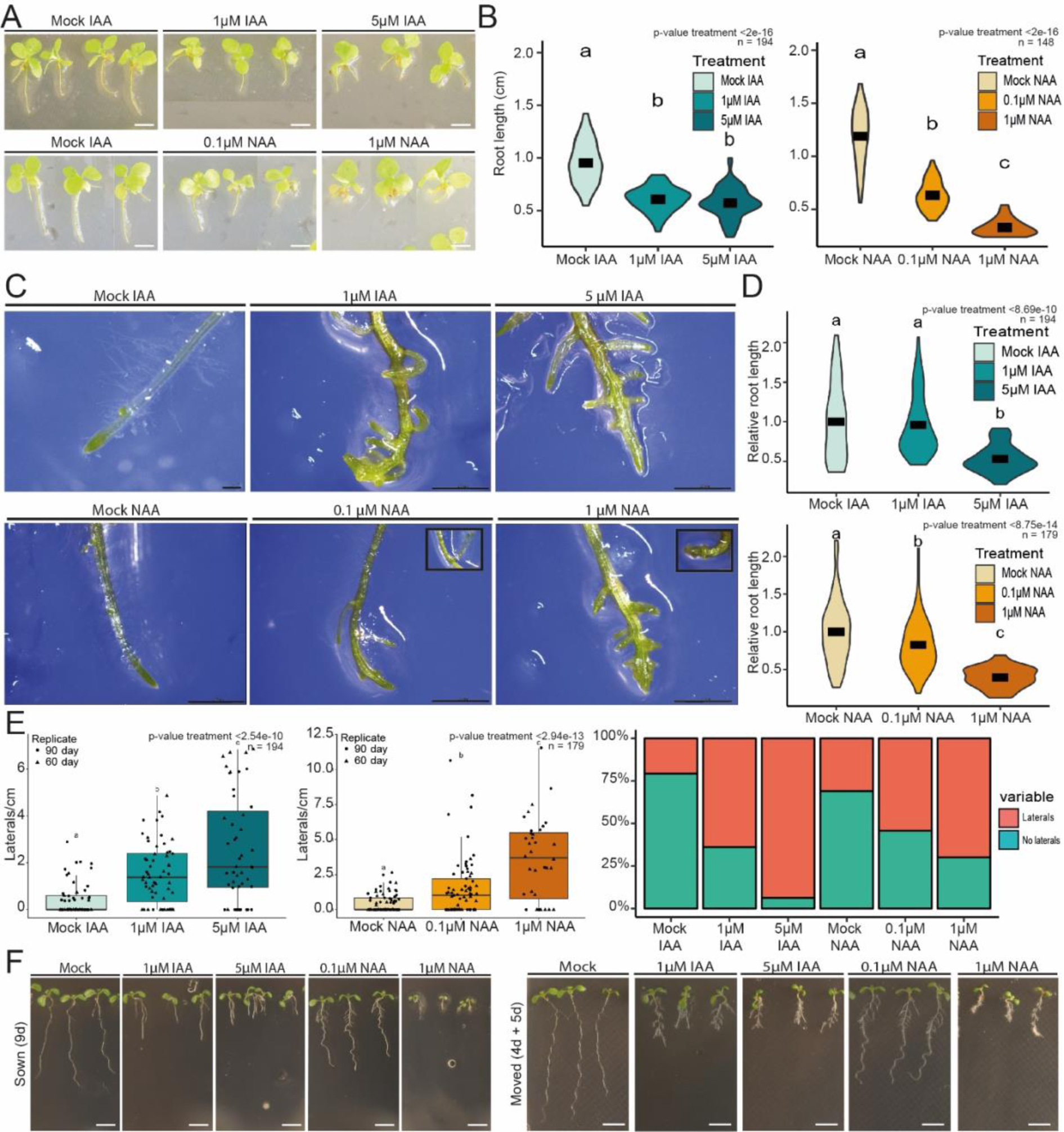
Auxin responses in Ceratopteris sporophytes. A) Young sporophytes grown on various concentrations of IAA or NAA for 12 days and B) Quantification of root length. C) Adventitious roots on sporophytes grown for 12 days in auxin-supplemented medium. D) Quantification of root length of adventitious roots grown on auxin with one representative replicate shown. E) Quantification of lateral roots per cm root and the frequency of roots bearing lateral roots under the different auxin conditions of a representative replicate. F) Representative pictures of Arabidopsis seedlings sown or transferred to auxin medium for the indicated times. Scale bars are 5 mm in A, 2.5 cm in C and 3 mm in F. Statistical significance shown by letters based on ANOVA and Tukey pair-wise comparison (P<0.05). ANOVA P-values plotted for treatments effects in plots.

The tissues that show responses to auxin in the Ceratopteris sporophyte and gametophyte are not homologous, except for root hairs, which are transcriptionally homologous to rhizoids (Jones and Dolan, 2012; Menand et al., 2007). We therefore analysed the response to auxin in root hairs. Root hairs showed an increase in length and appear to initiate closer to the root tip (Fig. S2B,C) in auxin-treated roots. Thus, Ceratopteris roots show a combination of sporophyte-specific responses that are similar to those in Arabidopsis (Fig. 2F), and a response shared with the gametophyte in both Ceratopteris and Marchantia.

We next explored leaf development, as it represents a laminar morphology similar to the gametophyte. Notably, however, fern leaves are not direct homologs of angiosperm leaves, whereas their roots are regarded truly homologous structures (Pires and Dolan, 2012; Szövényi et al., 2019; Tomescu, 2009). Upon IAA treatment, no clear difference in leaf shape was visible, but inhibiting auxin export (e.g., with NPA) resulted in highly malformed leaves with altered patterns of vascular tissues, which partly resembles NPA-treated Arabidopsis leaves (Mattsson et al., 1999; Verna et al., 2015) (Fig. S3). NPA-grown Ceratopteris leaves had higher cardinality and connectivity indices than control leaves, suggesting that auxin transport inhibition leads to the formation of more veins that are more frequently connected and therefore that efficient auxin transport inhibits vein formation and connection. The effect of auxin transport inhibition was particularly striking in the primary embryonic leaf: Whereas in control embryonic leaves the two veins branching off the single midvein remained unconnected, in NPA-grown embryonic leaves the vein branches connected into a loop. Together, those observation suggest a conserved role for auxin transport in vein formation of megaphylls.

### Disrupting auxin transport can induce a sporophytic program

When exploring the effect of auxin transport inhibition in gametophytes (Fig. 1F), we noticed that, in addition to the abnormal morphologies and increased rhizoid production, tissues also initiated root-like structures after prolonged treatment. These structures formed on gametophytic thallus and produced root hairs (Fig. 3A). The anatomy of these structures was indistinguishable that of sporophytic roots (Fig. 3D) Additionally, these roots produced adventitious vascular-like tissue in the gametophyte, recognizable by the distinct spiraling secondary cell wall thickenings (Fig. 3B). Importantly, these roots did not originate from archegonia (Fig. 3C), such as is the case for the apomictic *Dryopteris affinis* (Ojosnegros et al., 2024). To verify the ploidy of the gametophyte-derived roots, a transgenic H2B-GFP expressing lines (Geng et al., 2022) was imaged and the intensity of GFP fluorescence (Fig. 3E) was quantified as a proxy for ploidy. Sporophyte leaves and gametophytic thallus showed clearly distinct fluorescence intensities, consistent with their difference in ploidy (Fig. 3E), validating this approach for inferring ploidy. We found that sporophytic roots and NPA-induced gametophyte-derived roots showed no clear difference in GFP levels, indicating a similar (i.e. 2N) ploidy (Fig. 3E), in line with an additional quantification of DAPI fluorescence (Fig. S4D). Expression levels of the endogenous gene whose promoter was used to drive H2B-GFP expression (pCrHAM (Geng et al., 2022)) were comparable between gametophyte and sporophyte tissues (Fig. S4B), suggesting that ploidy, not gene expression, creates the differences in H2B-GFP observed.

**Figure 3:**
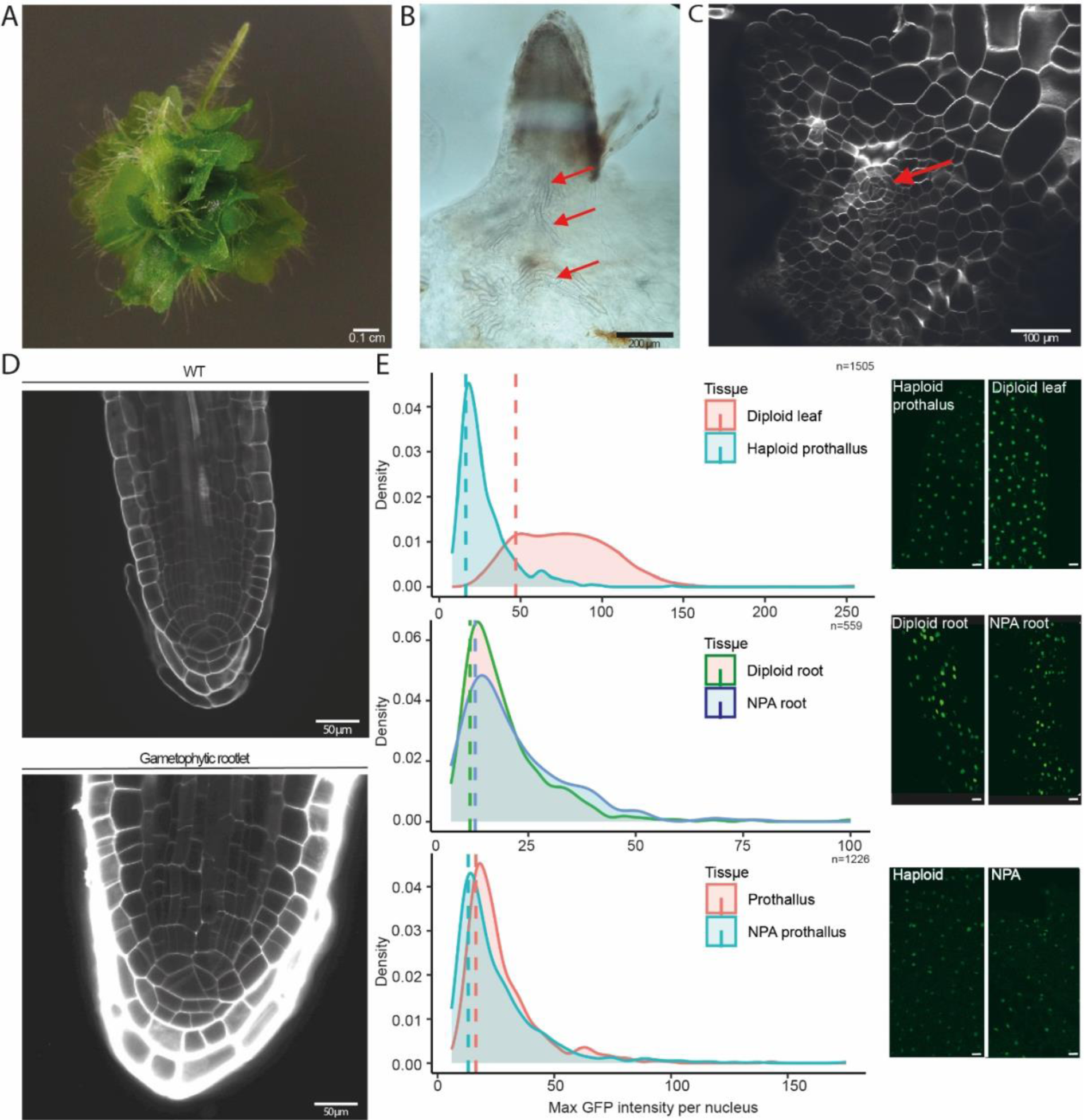
Auxin-induced ectopic root formation on gametophytes. A) Phenotype of gametophytes grown on unsupplemented media for 25 days after transfer from media containing 5uM NPA. B) Cleared ectopic root shows that vascular elements (red arrow) are formed in the gametophyte, that are not interconnected. C) Cleared gametophyte showing the induction of a root apical cell (arrow) on prothallus tissue. D) Comparison between sporophyte-derived roots and NPA-induced gametophyte-derived roots on a cellular level with both showing the distinct apical cell and root cap. E) Ploidy analysis by quantifying GFP fluorescence intensities in the root between haploid thallus and diploid leaves, diploid sporophytic roots and NPA-induced roots, and haploid thallus and NPA-grown gametophytic thallus. The right images show examples of GFP patterns in the two conditions for each comparison. Scale bars are 1 mm in A, 200 µm in B, 50 µm in C and 50 µm in D.

Consistent with NPA altering auxin distribution, NAA too could ectopically induce roots on gametophytic tissue (Fig. S4A). Upon scoring the frequency of root initiation (Fig. S4C), root initiation was most prevalent upon releasing plants from NPA treatment, going up to 50% of plants developing an ectopic root. Keeping the plants continuously on NPA also resulted in ectopic root formation but only after long cultivation times (50 days). Together, this suggests that disrupting auxin maxima triggers the initiation of root primordia, but outgrowth only happens upon restoring auxin transport. Interestingly, this root outgrowth only seems to happen upon transferring media and takes time, explaining why this response was not observed previously (Withers et al., 2023).

### Phylogeny and ontogeny conditions transcriptional auxin response

Transcriptional responses to auxin show clear differences between three bryophytes (*P. patens, M. polymorpha* and *A. agrestis*) and two vascular plants (*C. richardii* and *A. thaliana*) (Mutte et al., 2018). There are two main differences: 1) The ratio of auxin-activated versus auxin-repressed genes is shifted towards activation in vascular plants, and towards repression in bryophytes; 2) the amplitude of auxin-activated gene expression is higher in vascular plants. Given that sporophyte tissue was sampled for the vascular plants, and gametophyte tissue was sampled for the bryophytes, it is unclear which of these differences are due to phylogeny and which to tissue ontogeny.

We therefore compared transcriptional responses to auxin in both generations of Ceratopteris. We found that the amplitude of gene activation is higher in the sporophyte than in the gametophyte (Fig. 4A,B), suggesting that this difference is conditioned by ontogeny. The sporophytic response appears to be robust between experiments, as it overlapped significantly with prior published data (Fig S5D) (Mutte et al., 2018). By contrast, we found that in both generations, there is a dominance of gene activation, unlike in bryophytes (Fig 4A,B). This suggests that this trait is defined by phylogeny.

**Figure 4:**
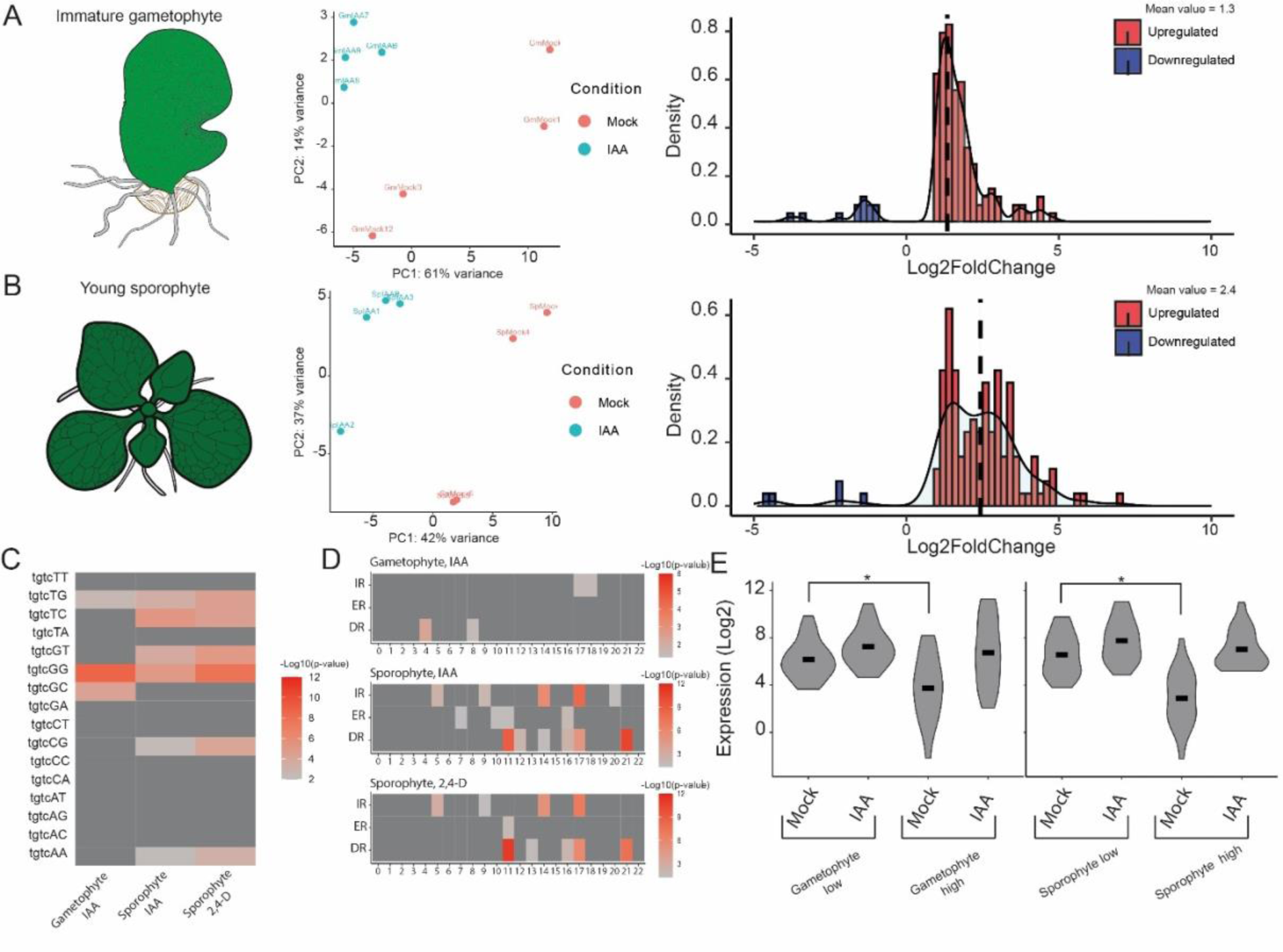
Transcriptional auxin response of Ceratopteris gametophytes and sporophytes. A,B) [left] Schematics of gametophyte (A) and sporophyte (B) stages on which the treatment was done [center] PCA plots showing the treatment effect and [right] density histograms of all DEG (fold change of IAA/mock). Average value is indicated with dashed line, upregulated genes are in red and downregulated genes in blue. C) The enrichment of 16 TGTCNN hexamers in auxin responsive promoters for the two generations of differentially expressed genes from Ceratopteris. Color denotes the significance of enrichment, -Log10(p-value). D) Abundance of TGTCNN repeats in auxin responsive promoters of differentially expressed genes from Ceratopteris. X axis shows the spacer length, Y axis denotes the fraction of genes with the repeats of specific structure. E) Violin plots showing the RNA expression values (Log_2_(TPM)) for the 20 lowest and 20 highest DEG in the gametophyte (left) and sporophyte (right). *: Wilcoxon-rank test P-value<0.005.

A promoter analysis on the differentially expressed genes showed conserved enrichment of the high-affinity ARF binding AuxRE (TGTCGG) in both generations (Fig. 4C). ARFs bind AuxRE repeats cooperatively as homodimers (Boer et al., 2014; Korasick et al., 2014; Nanao et al., 2014), and we explored the syntax of such repeats in auxin-regulated genes in both generations. This revealed differential enrichment of spacing in tandem repeat motifs (Fig. 4D). The sporophyte motif closely resembled those motifs that were previously reported in Arabidopsis and maize (reviewed in (Rienstra et al., 2023)). By contrast, the gametophyte-enriched motifs were weakly enriched, but distinct from the sporophyte-enriched ones. This suggests that the targets are indeed discrete and might differ in their mode of activation and that there is a sporophytic-tracheophyte specific motif spacing.

As is evident from the differential motif enrichment, there was only a small overlap in the genes controlled by auxin between the two generations (Fig. S5E). This overlap consisted of well-known auxin-inducible genes including *Aux/IAA*’s, *GH3*’s, YUCCA and *EXPANSIN*’s (Fig. S5A-C). Upon ortholog grouping and comparison, these genes are shared both with Marchantia and Arabidopsis. However, there was no clear enrichment of overlap between Arabidopsis and Ceratopteris sporophytes, or Marchantia and Ceratopteris gametophytes (Fig. S5A-C). Thus, we could not define clear shared sporophytic or gametophytic auxin-regulated genetic programs.

The amplitude of gene activation is defined by expression level in the absence and presence of auxin. High amplitude can therefore be generated by efficient repression in the absence of auxin, or by effective activation in its presence. We explored the likely mechanism underlying the difference in amplitude between gametophyte and sporophyte. The difference between highly and lowly upregulated genes was clearly due to an increase in repression in both generations (Fig. 4E). By comparing the two generations, we found that the 20 most strongly activated genes in sporophytes show a lower expression (log2 = 2.76) under mock conditions than in gametophytes (log2 = 3.78), whereas their expression upon auxin treatment was more similar between generations (log2 = 6.75 vs log2 = 7.34). This suggests that the increased amplitude of gene activation in the sporophyte is caused by efficient repression.

### Aux/IAA expression levels condition auxin sensitivity

We found that Ceratopteris gametophytes respond to auxin in a manner similar to the Marchantia gametophyte. Transcriptional responses are a direct outcome of the components of the Nuclear Auxin Pathway (NAP) encoded by each species’ genome, and expressed in each tissue. To identify possible genetic drivers of differences between gametophyte and sporophyte auxin response, we explored expression patterns of NAP components. The core components are: (1) the DNA-binding Auxin Response Factors (ARF), divided in auxin-dependent activating (A-class) and auxin-independent, repressing (B-class) (2) Aux/IAA repressors and (3) TIR1/AFBa that promote Aux/IAAs degradation in the presence of auxin (Lavy and Estelle, 2016). We found that *Aux/IAA* genes are more prominently expressed in the Ceratopteris sporophyte (Fig. S6). This analysis was further extended to publicly available transcriptomes from Ceratopteris across more developmental stages (Marchant et al., 2022; Marchant et al., 2019), showing a similar pattern (Fig. S7). This is consistent with an increased ability to repress target genes in the absence of auxin. However, A-class ARFs also seemed more abundant in the sporophyte generation, possibly also explaining the difference in response. B-class ARFs showed increased expression in the gametophyte, but expression levels were generally extremely low, and it is unclear what the biological significance of changes in expression at such low expression levels is.

Our transcriptomic data, along with gene content and differential expression of NAP components support a model where differential Aux/IAA expression and/or A-class ARF expression between generations creates distinct potential for high-amplitude gene activation. Indeed, relative to Marchantia, Ceratopteris has undergone duplications both in the Aux/IAA family and the A-class ARF subfamily (Mutte et al., 2018). To directly test the contribution of A-ARF or Aux/IAA copy number to auxin responsiveness, we generated transgenic Marchantia lines that either had slightly increased expression (Log2 = 4) of a mCitrine-tagged copy of MpARF1 (A-class ARF), or that express an additional copy of MpIAA, under the MpARF2 or MpARF1 promoter. Increased MpARF1 expression was achieved by complementing the *arf1-4* loss of function mutant (Kato et al., 2017) with a tagged, transgenic copy of the wild-type protein, and by selecting a line that had increased *ARF1* expression compared to wild-type plants (Fig. S8B). The advantage of Marchantia is the relatively simple system of only one A-class ARF and one Aux/IAA (Flores-Sandoval et al., 2015; Kato et al., 2020). Transgenic plants were tested for their auxin responsiveness by measuring thallus growth inhibition by a range of auxin concentrations (Fig. 5A-D). Increased MpARF1 levels did not induce a clear difference in auxin response compared to Tak-1 wildtype (Fig. 5B). By contrast, lines expressing an additional copy of MpIAA under the MpARF2 promoter showed a difference in overall phenotype and an increase in auxin sensitivity compared to Tak-1 (Fig. 5D). Like the endogenous MpIAA protein (Das et al., 2022), the additional copy was undetectable in control treatments in transgenic lines, but accumulated upon inhibition of the proteosome (Fig. S8A).

**Figure 5:**
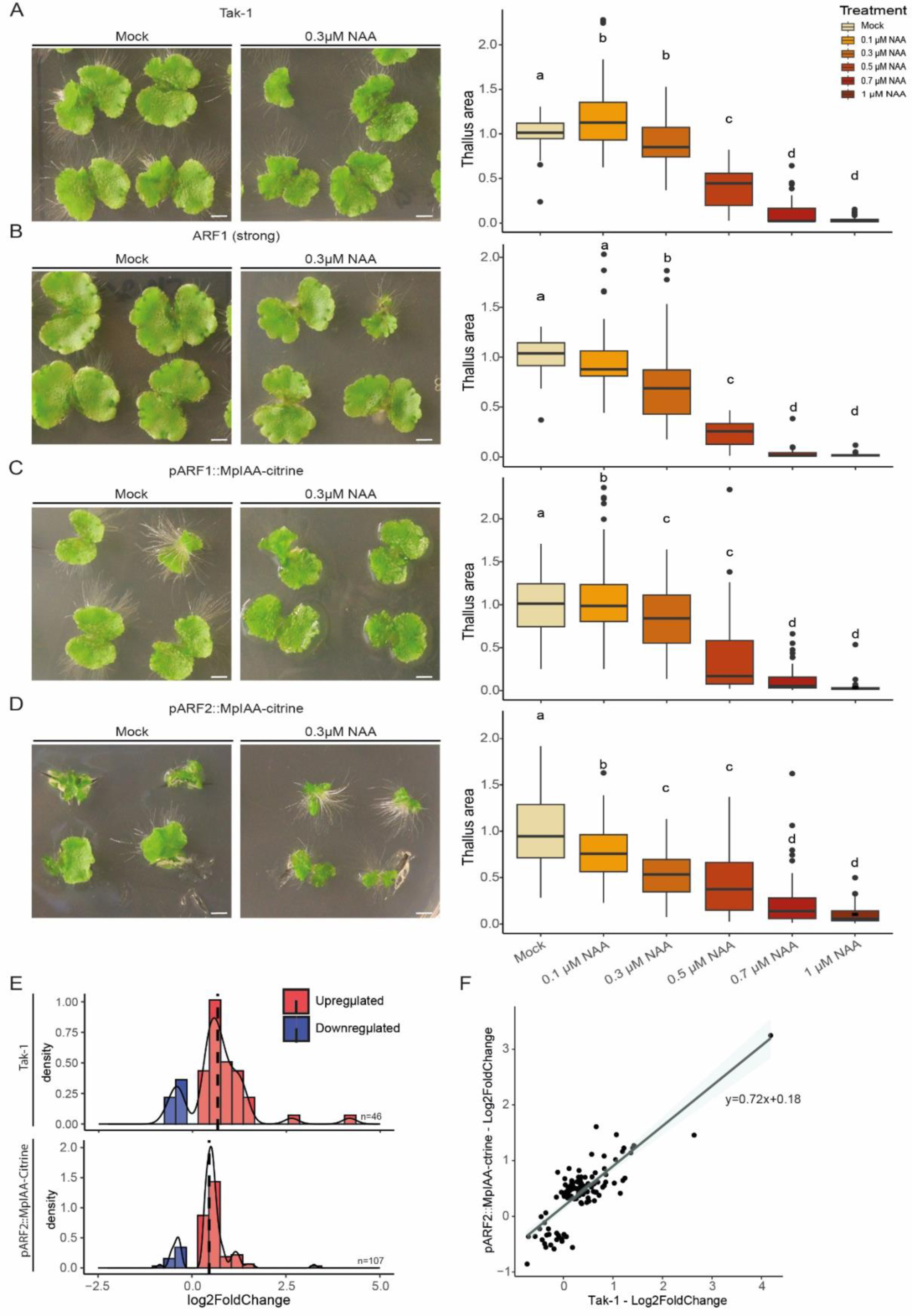
modulating auxin response characteristics in Marchantia. *A-D) Tak-1 wild type (A), pARF1-ARF1 (B), pARF1-MpIAA (= relative weak expression) (C) and pARF2-MpIAA (= relative strong expression) (D) gemmalings grown on mock media and 0.3 µM NAA (left panels), and quantification of relative projected thallus area of three combined replicates for each of the genotypes and across a range of NAA concentrations (right panels). E)). F) RNA-seq of* Marchantia *pARF2:MpIAA and Tak-1 density histograms of auxin responsive genes (P_adj_<0.05, Log_2_Foldchange(>0.3) F) Expression values of all auxin-responsive genes in either of two genotypes and the linear trendline describing the point cloud. Scale bars are 2.5 mm in A-D. Statistical significance shown by letters based on ANOVA and Tukey pair-wise comparison (P<0.05)*.

The increased auxin responsiveness in lines expressing an addition Aux/IAA copy is consistent with our predictions, but may or may not reflect a higher amplitude in auxin-dependent gene activation. To directly test the properties of gene expression output, we performed RNA-seq upon auxin treatment on a representative *proMpARF2-MpIAA-Citrine* line. We did not detect a clear shift in response amplitude of gene activation between Tak-1 and *proMpARF2-MpIAA-Citrine* plants (Fig. 5E,F). We used non-stringent filtering of the differentially expressed genes (Padj<0.05; Foldchange>1.25) to identify differentially expressed genes despite the low-inductive nature of auxin responses in Marchantia. With this filtering, a substantially larger group of genes is differentially expressed in the MpIAA-Citrine lines compared to Tak-1 (Fig. 5E), suggesting increased responsiveness. Upon plotting only the expression of genes that are auxin-responsive in the MpIAA-Citrine line, a slight increase of repression was visible (Log2(TPM)=8.93 instead of 9.25 in Tak-1). However, this was also accompanied by an increase in activation (9.7 vs. 9.37; Fig. S8C), showing a mixed response on a transcriptional level.

Based on these experiments in Marchantia, we conclude that Aux/IAA expression level does increase the degree of auxin-responsiveness, but that the sporophyte-like increased transcriptional response amplitude is likely caused by factors beyond Aux/IAA dose.

### Conserved rapid auxin responses in *Ceratopteris*

In addition to transcriptional responses, auxin promotes a number of fast cellular responses including cytoplasmic streaming and altered proton transport across the plasma membrane (Ayling and Clarkson, 1996; Barbez et al., 2017). We recently found that some of these responses are mediated by proteome-wide rapid auxin-triggered protein phosphorylation involving a conserved RAF-like protein kinase (Kuhn et al., 2024). Phosphoproteomic profiling in Arabidopsis, Physcomitrium, Marchantia and even streptophyte algae showed conservation of the response, and identified sets of common targets, as well as bryophyte-specific and species-specific targets. Again, given that sporophyte tissue was sampled for Arabidopsis, where as gametophytes were used for the two bryophyte species, it is hard to tell which of these differences are due to phylogeny, and which to ontogeny. We therefore performed phosphoproteome profiling upon 2 minutes of treatment with 100 nM of IAA in both gametophytes and sporophytes in Ceratopteris. Auxin triggered differential protein phosphorylation in both generations, but the number of auxin-regulated phosphor-targets was modest when compared with Arabidopsis or bryophyte species. This may in part be because we sampled whole sporophytes to make a fair comparison with the whole gametophyte, whereas for Arabidopsis, only roots were sampled. When comparing the two generations in Ceratopteris, we did not find clear differences in the profile of phosphorylation (Fig. 6A,B). Auxin-regulated phosphorylation in both generations more closely resembled that in bryophytes, than that in Arabidopsis. Both shared and unique functions are targeted by auxin-triggered phosphorylation in the two generations (Fig. 6A,B). Among these shared GO-terms (“plant organ development”, “carbon starvation”, “response to blue light”), at least one is conserved across all species that were previously tested (Kuhn, 2024) .

**Figure 6:**
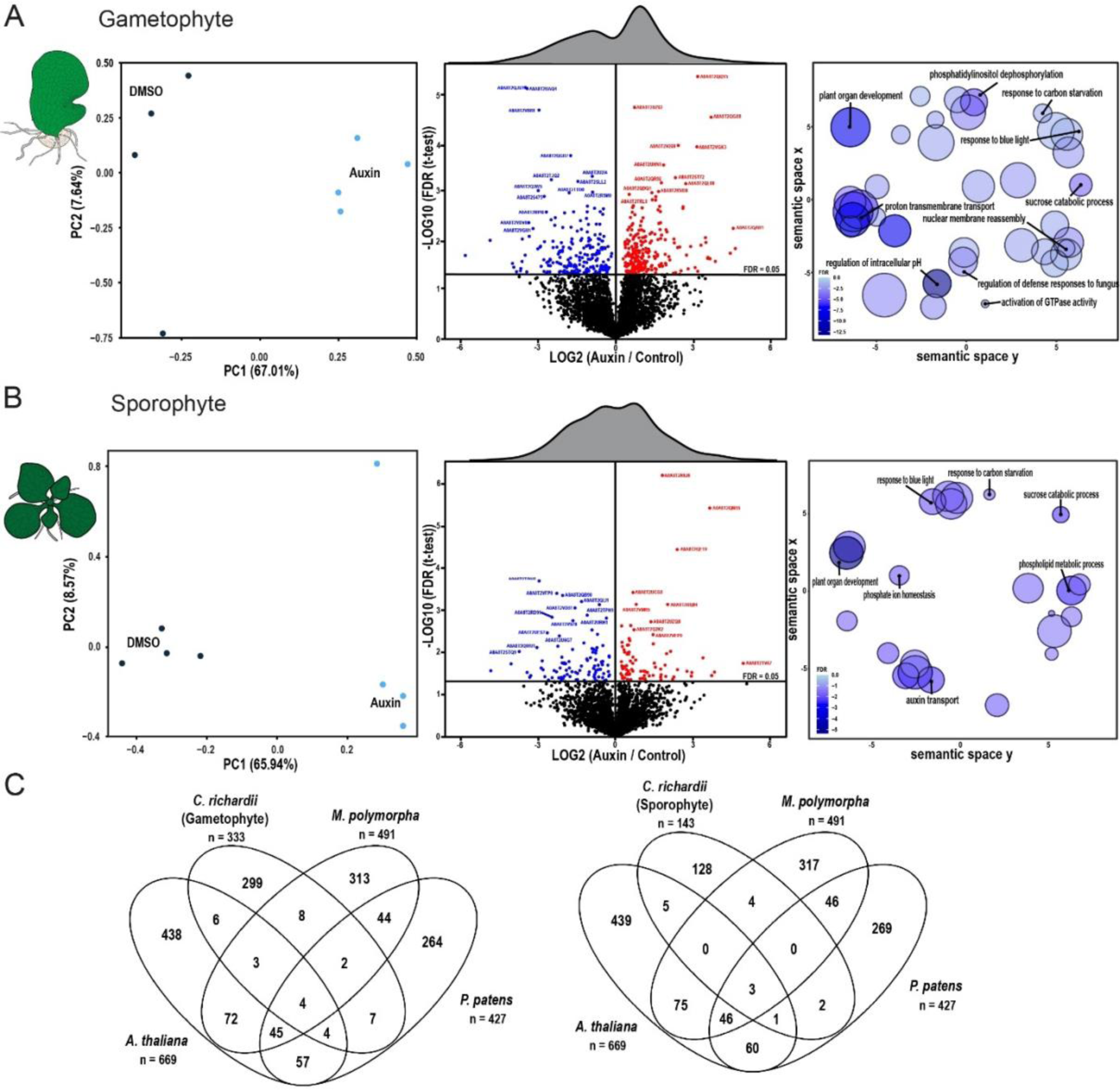
Auxin-triggered protein phosphorylation in gametophyte and sporophyte generations. *A,B) PCA analysis (left), differential phosphorylation (middle) and GO analysis of differentially phosphorylated proteins (right) upon 2 minutes of treatment with 10 nM IAA in Ceratopteris gametophytes (A) and sporophytes (B). C) Overlap of phosphosite orthogroups between the two Ceratopteris generations and previously reported datasets for* Marchantia *and Arabidopsis from Kuhn et al. (2024)*

We next explored the overlap between phosphor-targets between the two generations, and with Arabidopsis, Marchantia and Physcomitrium. In general, the overlap was limited (Fig. 6C) yet the number of overlapping terms is in the same order as between the five different species tested by Kuhn et al. (2024). This suggests that apart from a deeply conserved auxin-sensitive core, a wide range of species-specific and generation-specific changes are induced. Thus, auxin-triggered phosphorylation is conserved in Ceratopteris, and patterns of response are conditioned both by ontogeny and phylogeny.

## Discussion

Here we describe a set of different auxin responses in the model fern Ceratopteris across its two indeterminate, multicellular generations. On a phenotypic level, we see that gametophytes generally resemble thalloid bryophytes like Marchantia, while sporophytes mostly resemble flowering plants like Arabidopsis, in their capacity to respond to auxin. Auxin responses are numerous (Paque and Weijers, 2016), and it remains to be tested if the same patterns of response analogy hold for other growth or developmental processes. This similarity between Marchantia and Ceratopteris gametophytes is striking but is mirrored by the developmental homology with the same set of organs and cell types formed (i.e. rhizoids, antheridia, archegonia). One could even argue that Ceratopteris gametophytes are simpler than Marchantia gametophytes due to their short-lived nature and lack of a proper Z-axis development with no air chambers or storage tissue like in Marchantia (Conway and Di Stilio, 2020). Similarly, it appeals to reason that Ceratopteris sporophytes resemble *Arabidopsis* seedlings in their response to auxin, given that these species share the same evolutionary origin of their roots and vasculature (Szövényi et al., 2019). It is intriguing though that the response to auxin response inhibition in Ceratopteris leaves resembles that in Arabidopsis leaves, given that these leaves do not share an evolutionary origin (Pires and Dolan, 2012; Tomescu, 2009).

Long-term treatments with auxin or auxin transport inhibition in Ceratopteris gametophytes showed that sporophytic organs can be initiated, demonstrating the power of auxin as a developmental signal. To our knowledge, this is first report of such transdifferentiation, although it bears a superficial resemblance to the initiation of microspore-derived embryos in some flowering plant species (Corral-Martínez et al., 2020; Supena et al., 2008). It has previously been reported that “rod-like structures” develop from regenerating sporophytic callus in Ceratopteris (Xiao and Li, 2024). Likewise, roots are induced from eudicot callus when treated with a high ratio of auxin over cytokinin (Che et al., 2006). It has been reported that sugar in the growth media could induce apogamy in Ceratopteris gametophytes (Bui et al., 2017; Linh T et al., 2012). Since ectopic root formation did not depend on sugar, we interpret these structures as derived from a process distinct from apogamy. A plausible scenario is that auxin triggers reprogramming to a diploid state from which sporophytic organs can emerge. Identifying the intermediate stages and associated gene expression changes could help in identifying core root specification genes, as well as in the identifying sporophyte signature genes.

We also find that the amplitude of auxin response is stronger in the sporophyte than in the gametophyte, reflecting the bryophyte – tracheophyte split, and suggesting this to be an emergent property of ontogeny. It is unlikely that differences in tissue permeability to auxin contribute to these different responses, since Ceratopteris gametophytes are less complex in architecture, and most cells are directly exposed to the media. However, PINs (auxin efflux carriers) are less expressed in the gametophyte (not shown) and important for sporophytic development in Ceratopteris (Xiang and Li, 2024), which may dampen the response.

We tested directly whether the inferred duplications in A-class ARFs and Aux/IAAs that preceded the emergence of ferns may contribute to sporophyte-like response dynamics. Our results show that growth indeed becomes more sensitive to auxin when an additional Aux/IAA copy is expressed in the Marchantia gametophyte. However, gene expression does not fully resemble that in sporophytes of Ceratopteris or Arabidopsis in terms of amplitude. This result can be interpreted in many ways, but it is clear that there is additional complexity in the genetic architecture of the NAP and its differences between generations. A logical next step would be to combine the expression of an additional A-ARF copy with an additional Aux/IAA copy. Future advanced in facile genome editing in Ceratopteris (Jiang et al., 2024; Xiang and Li, 2024) and transgene expression (Geng et al., 2022; Plackett et al., 2014) will also help to further explore the genetic requirements for auxin response in both generations.

The above-mentioned similarities between Ceratopteris gametophytes and bryophytes suggest a conserved mechanism of auxin response based on their ontogeny. However, the targets of the auxin response are very divergent, besides the small core sets, as shown by orthogroup comparisons for both transcriptomics and phosphoproteomics. This is probably due to their distinct phylogenetic placement and 500 million years of divergence (Donoghue et al., 2021). Together with the previous points, this suggests that the exact targets of the response are strongly dependent on phylogeny, while the nature and mechanisms of the response depend more on ontogeny.

In summary, by studying the fern *Ceratopteris richardii*, a missing link in our understanding of auxin biology has been filled. The transcriptional nature of the response clearly depends on the ontogeny of the organism and vice-versa shapes its development due to their interdependency. This helps us to further understand the divergence, but equally importantly, the homology in hormone signalling between land plants. We expect that further exploration of the two generations in Ceratopteris will shed light on gametophyte and sporophyte developmental programs and their origin and homology.

## Acknowledgements

We are grateful to our team members and especially Sumanth Mutte and Danilo dos Santos Pereira for providing support with bioinformatic analysis and Aaron Ang for experimental support and Hirotaka Kato for providing the plasmids. This work was supported by a grant from the Graduate School Experimental Plant Sciences to S.W., a grant from the Human Frontiers Science Program (HFSP; contract number RGP0015/2022) to D.W., and grants from the Netherlands Organization for Scientific Research (NWO) to A.K. (VI.VENI.212.003) and J.R. (GSGT.GSGT.2018.013).

## Materials and Methods

### Plant growth conditions

Spores of *Ceratopteris richardii* strain Hn-n (Hickok et al., 1995) were sterilized and grown as described (Plackett et al., 2014) in a Hettich MPC600 plant growth incubator set at 28°C, with 16 hours of 100 μmol m^−2^ s^−1^. Plants were grown on ½-strength MS medium supplemented with 1% sucrose unless stated otherwise. Gametophytes were grown from spores and synchronized by imbibing the spores in the dark in water for >4 days. Sporophytes were obtained by flooding plates containing sexually mature gametophytes with water. *Marchantia polymorpha* plants were grown on ½-strength Gamborg’s B5 medium at 22°C with constant light. *Arabidopsis thaliana* plants were grown on ½-strength MS with 1% sucrose at 20°C, 60% humidity with 16h of light/day.

### Auxin growth experiments

For gametophytes, spores were sown directly on plates supplemented with Indole-3-acetic acid (IAA; Alfa aeser) or 1-Naphthaleneacetic Acid (NAA, Sigma). Size measurements were done when plants reached sexual maturity (±6/7 days). Alternatively, germinated spores were transferred onto auxin-supplemented medium before the lateral notch meristem was established (± 5 days) and grown for another 5 days. N-1-naphthylphthalamic acid (NPA, Sigma) treatments were done in a similar fashion. Gametophytes were imaged with a Leica M205FA epifluorescence microscope and their size was quantified by measuring their length and width. Sporophytes were grown in liquid 1/2MS without sucrose for root phenotyping. For every individual experiment, young sporophytes from the same plate were used to synchronize as much as possible their development. Sporophytes were grown for 12 days to measure root growth and branching. Sporophytes were imaged with a Canon EFS (18-135mm) camera, close-up of the root tips was done with a Leica M205FA epifluorescence microscope. Root lengths were measured using ImageJ and scored manually for the number of lateral roots. Rhizoid images were taken 3 days after transfer. Marchantia and Arabidopsis plants were treated in a similar manner, instead that the medium was different as described in the previous section.

To induce sporophytic roots on gametophytic thallus, spores were grown for at least 10 days on ½ strength MS containing 1% sucrose and 5 µM NPA, afterwards, they were transferred to NPA-free medium. First roots appeared 25 days after transfer. Sporophytic roots were also obtained when gametophytes were grown for 50 days on 5 µM NPA or if germinated normally and then transferred to 5-10 µM NPA or 5-10 µM NAA.

### Leaf venation analysis

NPA treated leaves were harvested 2 weeks after young sporophytes were transferred to medium containing 10 or 20 µM NPA. Only the youngest developed leaves were harvested to ensure that leaf primordia developed under NPA conditions. Leaves were fixed and cleared in ethanol: acetic acid for at least 1 day. Afterwards, leaves were rehydrated in 70% ethanol and stored at 4°C until imaging. Imaging was done with a Leica M205FA epifluorescence microscope. Quantification was done similar to Verna et al. (2015). Briefly, the number of touch, end, break and exit points of the veins were calculated for every leaf. From those numbers, the connectivity, absolute cardinality, and continuity index were calculated. The relative indexes were used to enable the pooling of different experiments. Statistical analysis was done by first testing for equal variance with a F-test and then a student’s t-test to test for equal means.

### Ploidy analysis

CrHAM::H2B-GFP line spores were described in Geng et al. (2022) and grown similarly as described before. GFP intensities were imaged with a Leica SP5 confocal microscope, with HyD detectors on photon counting mode to facilitate quantification. Z-stacks were obtained throughout the whole tissue, and maximum projections of those stacks were subsequently quantified in ImageJ. Nuclear intensity was quantified by first selecting the nuclei by binarizing the image. By subsequently analyzing the particles, all nuclear ROIs could be selected which were then imported to the original image to measure their intensities. All ROIs smaller than a given size (arbitrary area<30) were discarded as they were outside the expected size range for nuclei. DAPI quantification was done in a similar fashion by staining cleared roots (Kurihara et al., 2021), overnight with 50 µg/µL DAPI and washed once before imaging.

### RNA isolation and sequencing

For RNA isolation, immature gametophytes were grown for 5 days on a 100 µm nylon mesh. Young sporophytes of 22 days after fertilization were transferred to a new plate with a 100 µm nylon mesh and collected 5 days later after which new roots had developed. At this stage, sporophytes had approximately 5 leaves. Auxin treatments were performed by dissolving an IAA stock in liquid ½-strength MS with no sugar to a final concentration of 1 µM IAA and a DMSO control. The medium was preheated to 28°C to prevent any cold shock on the plants. Plates were taken out of the incubator, flooded with IAA or DMSO, and returned to the incubator for 1 hour.

RNA was isolated with the QIAGEN RNeasy kit and Total RNA was treated with RNase-free DNase I set (QIAGEN). RNAseq libraries were prepared and up to 20 million 150bp paired-end sequences were collected by Illumina-sequencing by Novogene (Uk). RNA quality was checked using FastQC (www.bioinformatics.babraham.ac.uk/projects/fastqc) and reads were mapped by Salmon and the obtained raw read counts were normalized and differentially expressed genes (p_adj_<0.05) were identified using DEseq2. All plots were made with ggplot2 (https://cran.r-project.org/web/packages/ggplot2/index.html) besides the upset plot with UpSetR (https://cran.r-project.org/web/packages/UpSetR/index.html)

*For* Marchantia, gemmae were grown for 9 days at 22°C. IAA treatments were done as described previously by Kuhn et al. (2024). Briefly, prior to IAA treatment, plants were flooded with liquid medium overnight before incubating with 1 µM IAA for 1h. Marchantia RNA was isolated with the QIAGEN RNeasy kit with an additional Trizol step before column binding. RNAseq libraries were prepared and up to20 million 150bp paired-end sequences were collected by Illumina-sequencing by BMKGENE (Germany). RNA quality was checked using FastQC (www.bioinformatics.babraham.ac.uk/projects/fastqc) and reads were mapped by Hisat2 and the obtained raw read counts were normalized and differentially expressed genes (p_adj_<0.05) were identified using DEseq2.

### Expression analysis

For the comparison between gametophytes and sporophytes, the normalized expression values of the mock treatments of both life stages were compared using R package ComplexHeatmap (https://bioconductor.org/packages/release/bioc/html/ComplexHeatmap.html). Expression values across all developmental stages were retrieved from (Marchant et al., 2022) and TPM values were normalized to a Z-score which were plotted with ComplexHeatmap.

### Promoter analysis

We used the reference genome of fern *Ceratopteris richardii* v2.1 from Phytozome DB. Promoters of all genes were analysed for overrepresentation of hexamers in the interval 600 bp upstream from the transcription start site (TSS) to either translation start site or 1 kbp downstream TSS, depending on which was smaller. We took into analysis three sets of up-regulated DEGs (foreground sequences), and generated for each of them the background consisting of the same genomic intervals for the rest genes. First, for each pair ‘foreground vs. background’ sets, we used Fisher’s exact test to estimate the enrichment for TGTCNN consensus sequences. Here and below, this test counted the number of genes. Second, we applied the package MCOT (Levitsky et al., 2019) to count the numbers of genes containing in the genomic interval TGTCNN repeats with spacers from 0 bp to 25 bp, and specific orientations (direct (DR), inverted (IR) and everted (ER) repeats). For each pair ‘foreground vs. background’ and any possible mutual location and orientation of hexamer in pairs, we estimated the enrichment of their co-occurrence by Fisher’s exact test.

### Orthogroup analysis

Orthogroups between the different species were identified using Orthofinder (Emms and Kelly, 2019). Therefore, transcriptomes from the used species *Arabidopsis thaliana* (Araport11), *Marchantia polymorpha* (v6.1) and *Ceratopteris richardii* (v2.1) were used and common orthologous sequences were identified. Similarly, for the phospho-proteomics, orthogroups were identified from the proteomes of the different species. The auxin-responsive DEG or phosphosites from the different species were converted to their specific orthogroup and subsequently overlayed using a Venn diagram.

### Plasmid construction

MpARF1 and MpARF2 promoters (+ 3kb upstream) were amplified with the primer set HK120/HK125 and HK126/HK127 respectively (Table S1). and cloned into pMPGWB307 using the Xbal site (pHKDW031/038). MpIAA and MpARF1 CDS were subcloned into pENTR/D (thermo) using the primer combinations of MpIAA_entry/JHG081 and HK009/HK015 respectively. These genomic CDS sequences were then transferred to the pHKDW031 (pARF2) or 038 (pARF1) with Gateway LR Clonase II Enzyme mix (Thermo Fisher Scientific). pHKDW038 was described previously and kindly provided by Hirotaka Kato (Kato et al., 2017; Kato et al., 2020).

### RT-qPCR

MpARF1 expression was validated in multiple complemented arf1-4 mutant (Kato et al., 2017) to select for a higher expressing line. RNA was isolated from ten-day old gemmalings as described before. 1µg of total RNA was used for cDNA synthesis (iScript cDNA synthesis kit, Bio-Rad) according to manufacturer’s instructions. RT-qPCR was performed using a 384CFX Connect Real-Time PCR Detection system (Bio-Rad) and iQ SYBR Green Supermix (Bio-Rad). A two-step cycle of 95°C 10s followed by 60°C for 30s was repeated for 40 cycles, followed by a melt-curve analysis. Three biological and two technical replicates were used. All primers used are listed in Supplementary table 2. The geometric mean of MpSAND, MpAPT7 and MpAPT3 (Saint-Marcoux et al., 2015) was used to normalize expression of MpARF1 according to (Taylor et al., 2019).

### *Marchantia* transformation

A protocol based on the *Agrobacterium*-mediated transformation of *M. polymorpha* regenerating thalli (Kubota et al., 2013) was used. Briefly, Agrobacterium cultures were grown for 2 days in liquid LB medium. Afterwards they were spun down and resuspended in liquid Gamborg medium supplemented with sucrose and casamino acids and acetosyringone and left for 6h. Tak-1 thallus was cut into 1mm x 1mm pieces and added to liquid medium together with Agrobacterium. Co-cultures were grown for three days at 22 degrees while shaking. After washing, positive transformants were selected on medium containing chlorsulfuron (0.5uM) and cefotaxime (100mg/L). Transformants were validated by PCR and microscopy. The strong expressing line for ARF1 was validated by qPCR

### Phosphoproteomics

Ceratopteris gametophytes and sporophytes were grown on mesh and treated for 2 minutes with 100nM IAA and harvested immediately. Protein purification and phospho-peptide enrichment and measurements were done as described earlier (Kuhn et al., 2024).

### Confocal microscopy

Ceratopteris gametophytes grown on auxin-supplemented medium were cleared and fixed using Clearsee alpha (Kurihara et al., 2021), and stained with SR-2200/Renaissance (Musielak et al., 2016). Roots were cleared and stained in a similar fashion. Imaging was done with a Leica SP5 confocal microscope.

Marchantia MpIAA-Citrine was detected with a Leica SP8 confocal microscope. MG132 (Sigma) treatments were done on gemmae that were allowed to germinate for 8h in the presence of MG132 and imaged afterwards.

